# Challenging popular belief, mosquito larvae breathe underwater

**DOI:** 10.1101/2023.03.06.531304

**Authors:** Agustin Alvarez-Costa, Maria Soledad Leonardi, Silvère Giraud, Pablo E. Schilman, Claudio R. Lazzari

## Abstract

It is taken for granted that immature mosquito only breathe atmospheric air through their siphons. However, there is no quantitative study that demonstrated it. We analysed the survival of the last instar larvae of *Aedes aegypti* fully submerged at different temperatures, and measured oxygen consumption from air and dissolved in water, of larvae and pupae of this species under different conditions. Results revealed that under water, larvae survived much longer than expected, reaching 50% mortality only after 58, 10, and 5 days at 15°, 25° and 35°C, respectively. Interestingly, whereas we registered moults to pupae in larvae with access to air, individuals kept submerged never moulted. When remaining at the water surface, larvae obtained 12.72% of O_2_ from the water, while pupae only 5.32%. When completely submerged, larvae consumed less oxygen than in contact with the surface, but enough for surviving, while pupae did not. At both media, temperature affected larvae respiration rate, with relatively close Q_10_ values. In the related species, *Ae. albopictus*, a similar pattern of O_2_ consumption were observed. Larvae got 12.14% of their oxygen from the water. Interestingly, no significant differences in total O_2_ consumption were found between water O_2_ consumption, when *Ae. albopictus* larvae were submerged, or when they also have access to air (dual O_2_ consumption). Our findings not only challenge the classical idea that mosquito larvae only breathe atmospheric O_2_, but also force us to reconsider the potential effectiveness of control methods based on asphyxiating larvae by detaching from water surface.

**Summary statement:** We present the first quantitative analysis of mosquito larvae respiration in air and water, unravelling the unknown capacity of larvae of the most cosmopolitan disease vector of breathing underwater.

## Introduction

As an originally terrestrial group and from an evolutionary standpoint, when insects colonized the freshwater, have to deal with many vital functional problems. Respiration was probably the greatest one. As an environment, water is a less favourable medium for respiration because its oxygen concentration is twenty times lower, and its rate of oxygen diffusion is lower by a factor of 105 (Dejours, 1988). In this sense, insects that had adapted to water environments developed plenty of different strategies to acquire oxygen. Among them, there are examples of cutaneous respiration, developed spiracular, tracheal, and blood gills, the use of air stores carrying air bubbles, or the use of hydrofuge structures to remain attached to the water surface and breathe atmospheric air (Wigglesworth, 1972). It is taken for granted that the last is the case of mosquito larvae and pupae, who would use their siphons as “snorkels” when they rest attached to water surface. This statement is widely affirmed in textbooks, internet scientific and dissemination sites, and also in protocols for controlling mosquito populations at juvenile stages. Yet, the vital role of aerial respiration in immature mosquitoes has been challenged by sporadic observations since long time ago (e.g., Da Costa Lima, 1914; MacFie, 1917; Ramsey and Carpenter, 1932; Wang, 1938; Richards, 1941).

Even though it seems reasonable that atmospheric air would constitute the main source of oxygen, it cannot be excluded that mosquito larvae and pupae could gather some oxygen from the water. On the one hand, the survival underwater of larvae belonging to different mosquito species, along with some measurements of oxygen consumption from the water (Sen, 1914; Fraenkel and Herford, 1938) are pieces of evidence that water-dissolved oxygen could be sufficient for maintaining vital functions during periods without contact with the surface. However, the relative contribution atmospheric and water-dissolved oxygen to the metabolism of mosquito larvae and pupae has not been yet investigated.

In our study, we attempt to shed light on respiratory gas exchange in immature mosquito stages. We analysed the survival and oxygen intake of mosquito larvae and pupae having or not having access to air and measured the effect of the temperature on these variables. For the first time, we provide quantitative data on oxygen consumption from air and water, measuring them both, separately and simultaneously. Finally, we discuss the evidence obtained during our experiments in the framework of mosquito respiratory physiology, as well as the possible consequence of our findings for control methods based on the suffocation of juvenile mosquitoes

## Materials and methods

### Mosquitoes

Eggs of *Aedes aegypti* of the Bora strain (insecticide susceptible) and *Ae. albopictus* of Vectopole strain reared at Vectopole (Montpellier, France) were provided by the European network InFravec2 (https://infravec2.eu/). Eggs were put in dechlorinated tap water for hatching, adding traces of ascorbic acid and tropical fish food, and kept at 26°C (± 1°C) in a climatic chamber, under a light/dark cycle 12h:12h/ (lights on at 08:00 am). Food was regularly provided until they reached the 4^th^ instar (6-10 days in our conditions), and used for experiments. Individuals were handled by aspirating them with plastic Pasteur pipettes with their tip cut. Each larva was tested only once and discarded afterward.

### Survival experiments

The survival time of fourth-instar larvae of *Ae. aegypti* was evaluated at three temperatures (15°, 25°, and 35ºC). A climatic chamber was set at the experimental temperature, and these values were kept constant at nearly 1°C. A 12h:12h light cycle was imposed with lights on at 08:00 am. Relative humidity was kept at 70% to reduce the evaporation of the water in the recipients. In case some evaporation occurred, deionised water was daily added to return to the initial volume.

For each temperature, two larvae were placed in recipients with 300 ml of dechlorinated water under two conditions, submerged or with access to air. In both cases, a unique larva was placed into a glass cylinder (0.6 cm in diameter and 2 cm in length), both ends closed by a tissue mesh kept with the aid of an O-ring; to allow water circulation, but keeping the larva caged at the same time. In the first condition (submerged), the cylinder was completely sunk at the bottom of the recipient, taking care that no air remained captive inside. The control condition consisted of a larva placed in the same cylinder maintaining half of it in contact with the air, and the other half underwater (Fig. S1). No food was provided, but the water was neither changed, nor the development of microorganisms impeded. Recipients were placed inside a climatic chamber at 15°, 25°or 35°C (± 0.5°C), under a light/dark cycle at 12h:12h/ (lights on at 08:00 am). The number of dead and moulted insects was recorded daily.

### Oxygen consumption

The individual oxygen consumption of immature *Ae. aegypti* was measured using optodes. We employed two 4-channel Firesting O_2_ meter (Pyro Science, Aachen, Germany) using 4 ml vials with an integrated optical oxygen transducer (OXVIAL 4). Briefly, flashes of light of specific wavelengths generated in the interface are guided through a light fibre to excite a transducer inside the vial from the outside. The fluorescens of the substance, which is proportional to the oxygen concentration in the medium (air or water) is gathered by the same optic guide and analysed at the interface. The temperature of the vials was controlled using a Peltier element and controller (QuickCool 34W; Peltron Gmbh, Germany). A temperature sensor from the oxygen meter measured the temperature inside an empty vial, and its signal allowed the system to adjust the values of the measured O_2_ concentrations.

Fourth-instar larvae of *Ae. aegypti* and *Ae. albopictus*, and pupae of *Ae. aegypti* were evaluated. Most measurements were carried out on *Ae. aegypti* under three vial conditions: submerged, closed vial, and open vial, at 15°, 25° and 35°C. The condition submerged consisted in a closed vial filled with water with an individual inside, and water O_2_ concentration was registered for four hours. In this treatment, the total absence of air bubbles was carefully checked. For the closed vial condition, an individual of the mosquito immature stage was placed in a closed vial half filled with water, and the water and air O_2_ concentration was registered for four hours. Finally, the open vial condition consisted of an open half-filled water vial with an individual mosquito inside, and the water O_2_ concentration was registered for four hours. Each treatment was replicated between 12 and 30 times. The rate of oxygen consumption was calculated for each replicate, calculating the slope of a linear regression of the O_2_ concentration versus the time of the experiments. For *Ae. albopictus* same assays were performed except for open vial condition and close vials at 15 and 35°C.

### Statistics

To analyse differences in survival across temperatures and conditions (submerged and control) a Kaplan-Meier analysis was performed.

For the analyses of oxygen consumption, the effects of vial conditions (submerged, closed vial, and open vial), temperature (15°, 25° or 35°C), the medium where O_2_ was taken (air or water) and stage (larva or pupa) in O_2_ consumption one or two-ways ANOVAs were performed. *A posteriori* comparisons of significant ANOVAs were performed by means of Tukey test. The significance threshold was chosen at 0.05 for all analyses.

## Results

### Survival experiments

The survival of *Ae. aegypti* larvae were significantly affected by the temperature and immersion conditions (p<0.05). In the control treatment (i.e., with access to air) at 25°C, no death was registered, and the survival curve was significantly higher than the curve of the submerged treatment at the same temperature. At 35ºC, the survival curve of the submerged larvae presented the lowest values with the higher negative slope, differing significantly from its control at the same temperature. Surprisingly, at 15 ºC the submersion treatment did not differ from the control (Fig. 1A). The 50% mortality of submerged larvae differed with the temperature, being 58, 10, and 5 days at 15°, 25°, and 35ºC, respectively (Fig. 1B). Remarkably, some individuals remained alive for as long as 30 days at 25ºC and 68 days at 15ºC. Finally, whereas we registered moults to pupae in control larvae, individuals kept submerged never moulted (Table 1).

**Figure 1.**
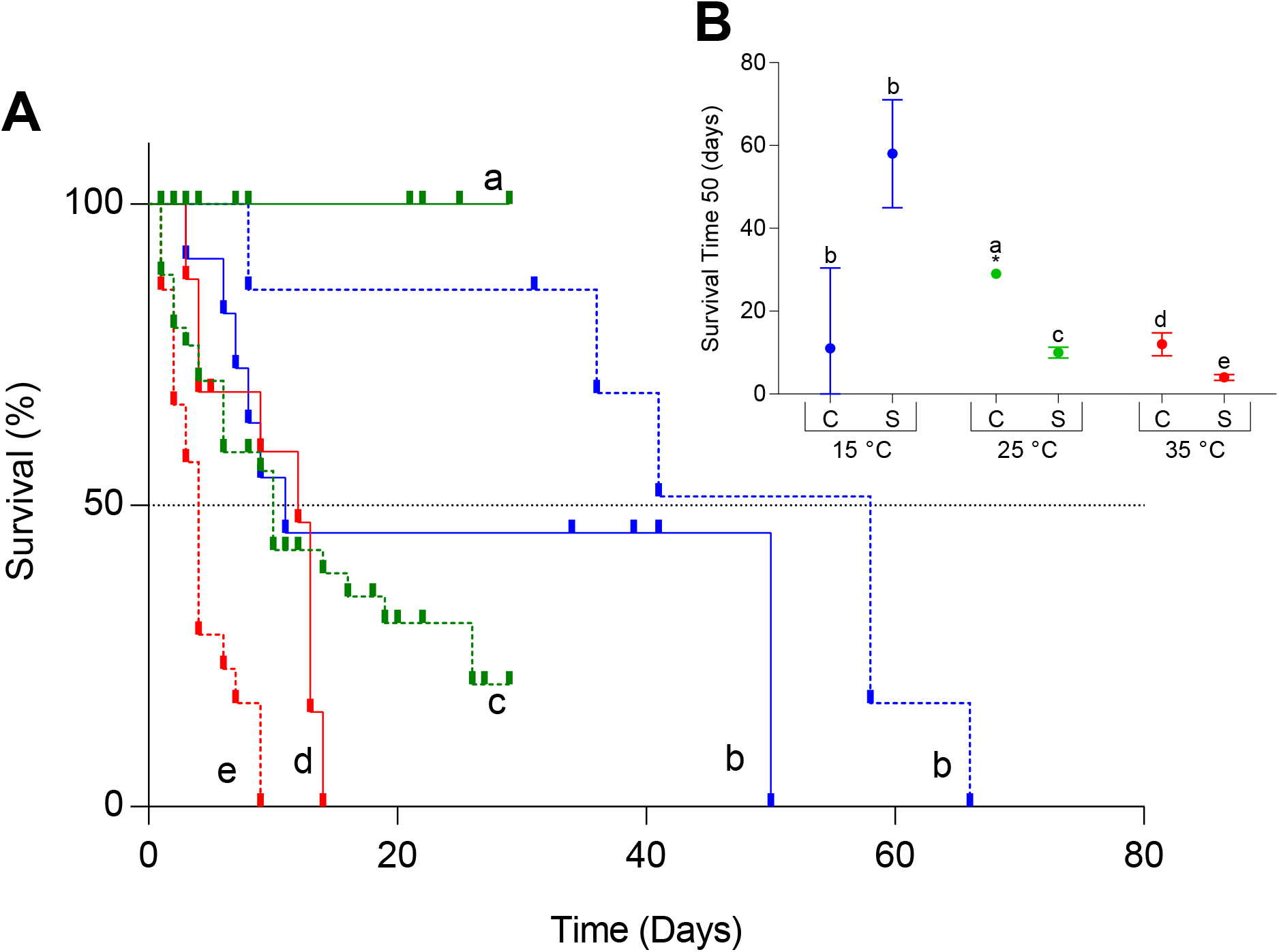
(**A**) Survival of *Ae. aegypti* larvae: control (solid line) and submerged treatments (dotted line) at three temperatures: 15°, 25°, and 35°C (blue, green, and red respectively). **(B)** Survival time 50 (estimated and 95 % confidence intervals) of *Ae. aegypti* larvae at control and submerged treatments at three temperatures: 15°, 25°, and 35 °C (blue, green, and red respectively). Different letters indicate significant differences between conditions (submerged and control) and between temperatures in the submerged condition (p<0.05). *It was not possible to estimate the survival time 50 due to the lack of mortality in the control treatment at 25 ºC, so the value shown is the maximum survival time registered.

**Table 1.**
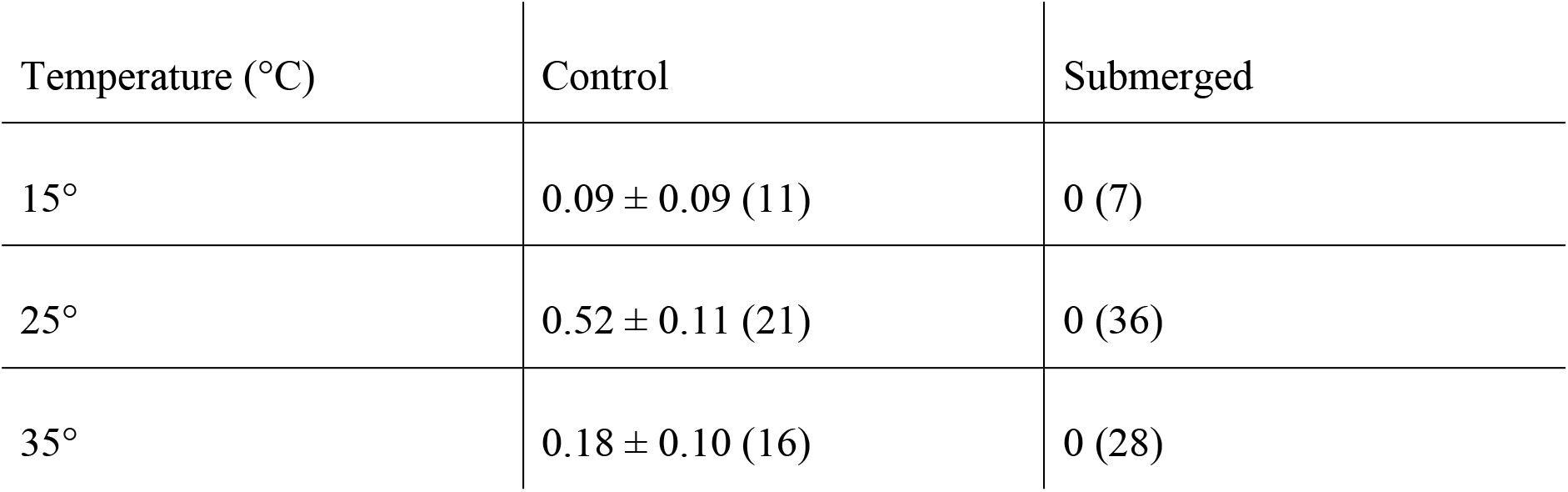
Proportion of pupae ± s.e.m. (number of individuals) of *Ae. aegypti* larvae at control and submerged treatments at three temperatures: 15°, 25°, and 35°C.

| Temperature (°C) | Control | Submerged |
| --- | --- | --- |
| 15° | 0.09 ± 0.09 (11) | 0 (7) |
| 25° | 0.52 ± 0.11 (21) | 0 (36) |
| 35° | 0.18 ± 0.10 (16) | 0 (28) |

### Oxygen consumption

#### Larvae and pupae at 25ºC

In all three conditions: submerged, closed vial and open vial, *Ae aegypti* larvae and pupae evinced to consume measurable amounts of oxygen from the water (Fig. 2). A significant difference in O_2_ consumption from the water was observed between the interaction of immature stages and the vial conditions (two-way ANOVA F: 107.31, DF: 5, p<0.0001). Larvae under the submerged condition presented the highest rate of water-dissolved O_2_ consumption, followed by pupae under the same vial conditions. In addition, larvae under closed vial and open vial conditions presented higher rates of O_2_ consumption than pupae under the same vial conditions, but their O_2_ consumption rates were always lower than larvae and pupa under submerged conditions (Fig. 2). No significant difference was found between closed and open vial treatments for both immature stages, allowing us to employ closed vials for next measurements.

**Figure 2.**
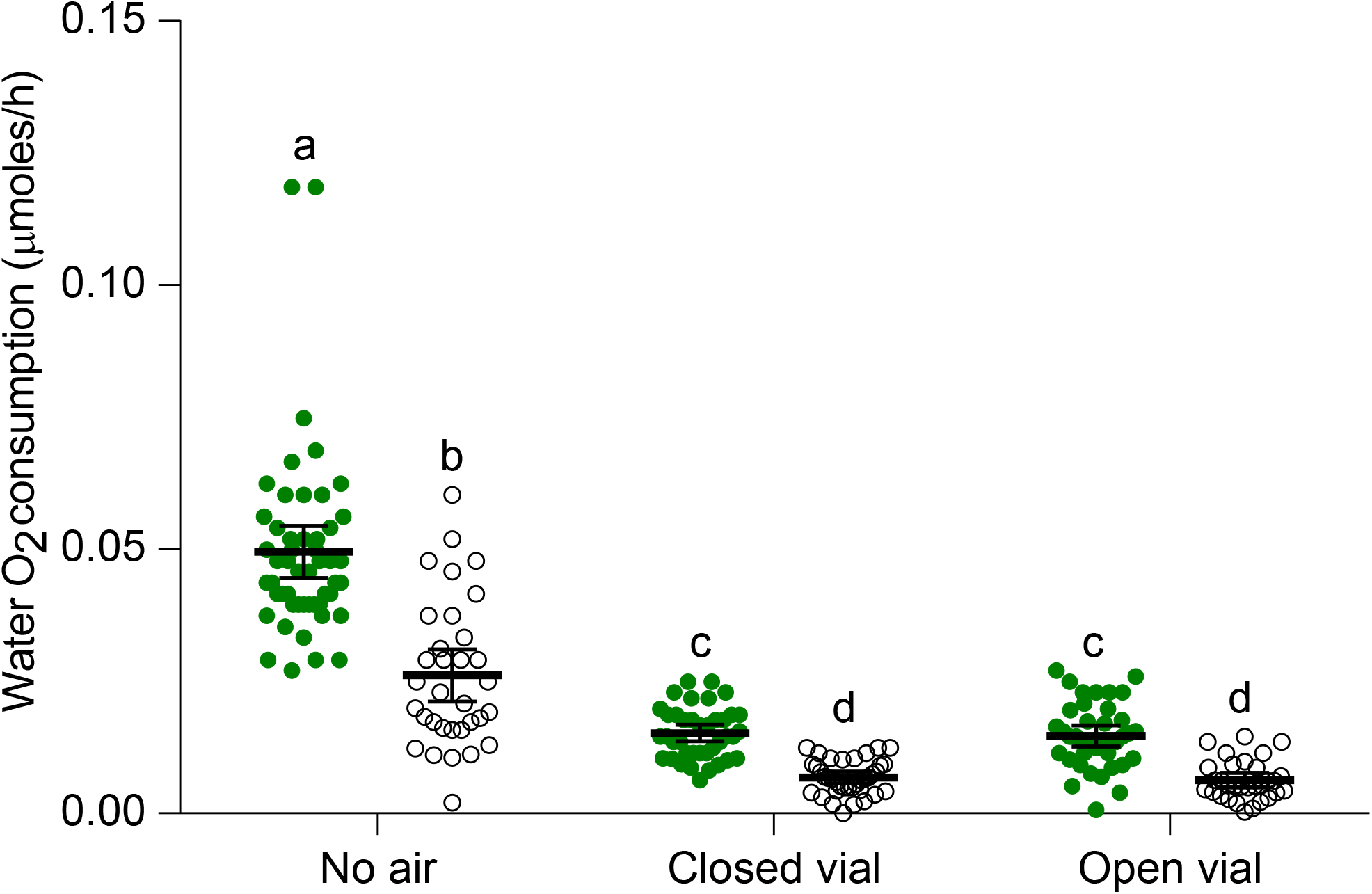
Consumption of water-dissolved O_2_ (mean and 95 % confidence intervals) by *Ae. aegypti* larvae and pupae under three conditions: submerged, closed vial and open vial. Different letters indicate significant differences between treatments (Tukey multiple comparison).

Pupae presented higher oxygen consumption from the air than larvae under closed vial conditions (two-way ANOVA, F: 119.75, DF: 3, p<0.0001, Fig. 3). The air O_2_ consumption of both immature stages tested was significantly higher than their water O_2_ consumption (p<0.05, Fig. 3). The dual rate of O_2_ consumption, adding air and water O_2_ consumption of closed vial treatment, were 0.111 and 0.128 moles/h for larvae and pupae respectively. Larvae obtained most of their oxygen from the air, but about 13% from water, while pupae obtained almost 95% from the air and the rest from water (Fig. 4B, C). Finally, the dual (air + water) O_2_ consumption in closed vial condition was significantly higher than O_2_ consumption from the water of the submerged condition (two-way ANOVA, F: 68.85, DF: 3, p<0.0001, Fig. 4).

**Figure 3.**
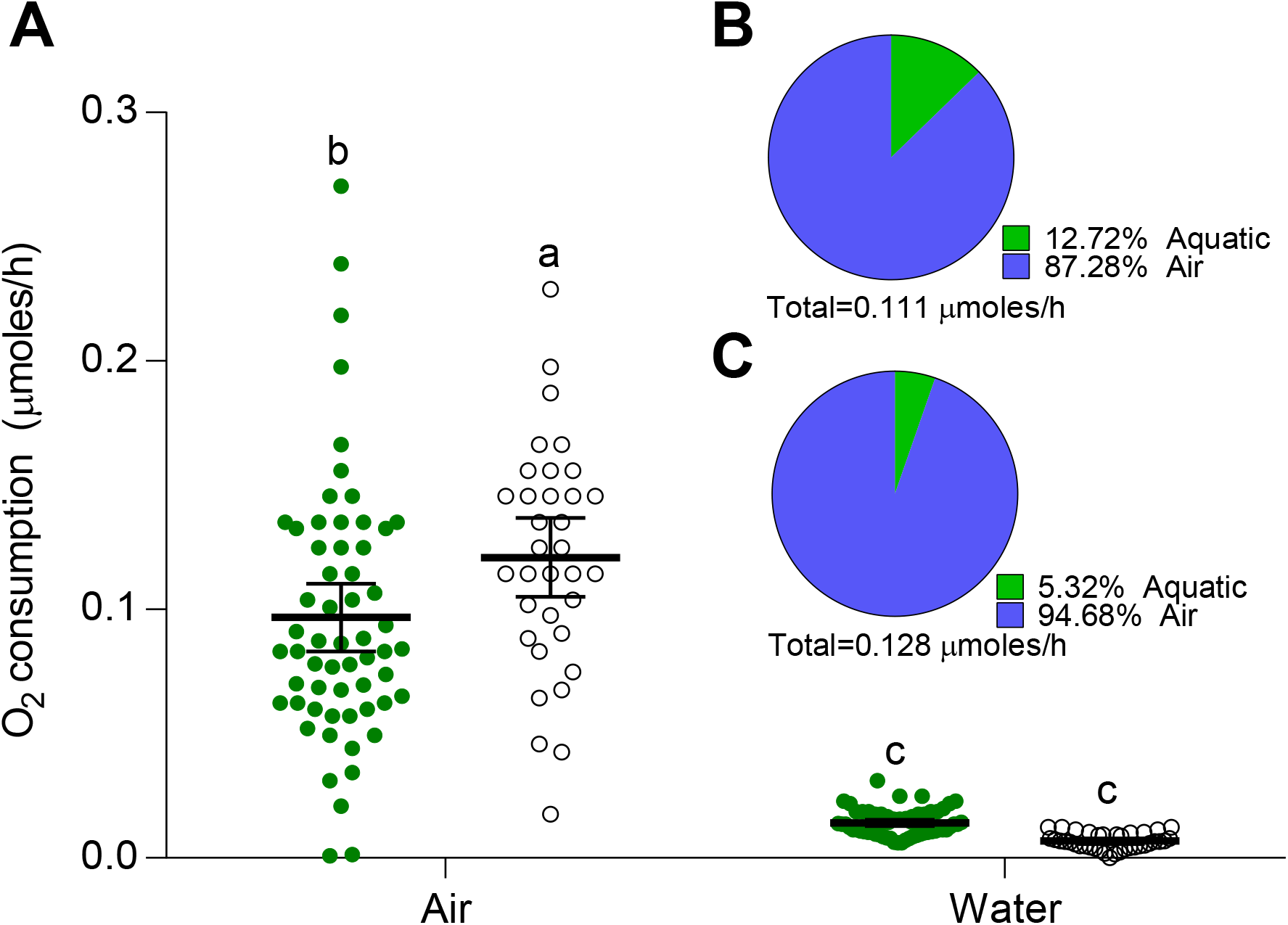
(**A**) O_2_ consumption (mean and 95 % confidence intervals) obtained from air and water of *Ae. aegypti* larvae and pupae in the closed vial condition. Different letters indicate significant differences between treatments (Tukey multiple comparison). Percentage of oxygen consumption of *Ae. aegypti* larvae (**B**) and pupae (**C**), obtained from air and water.

**Figure 4.**
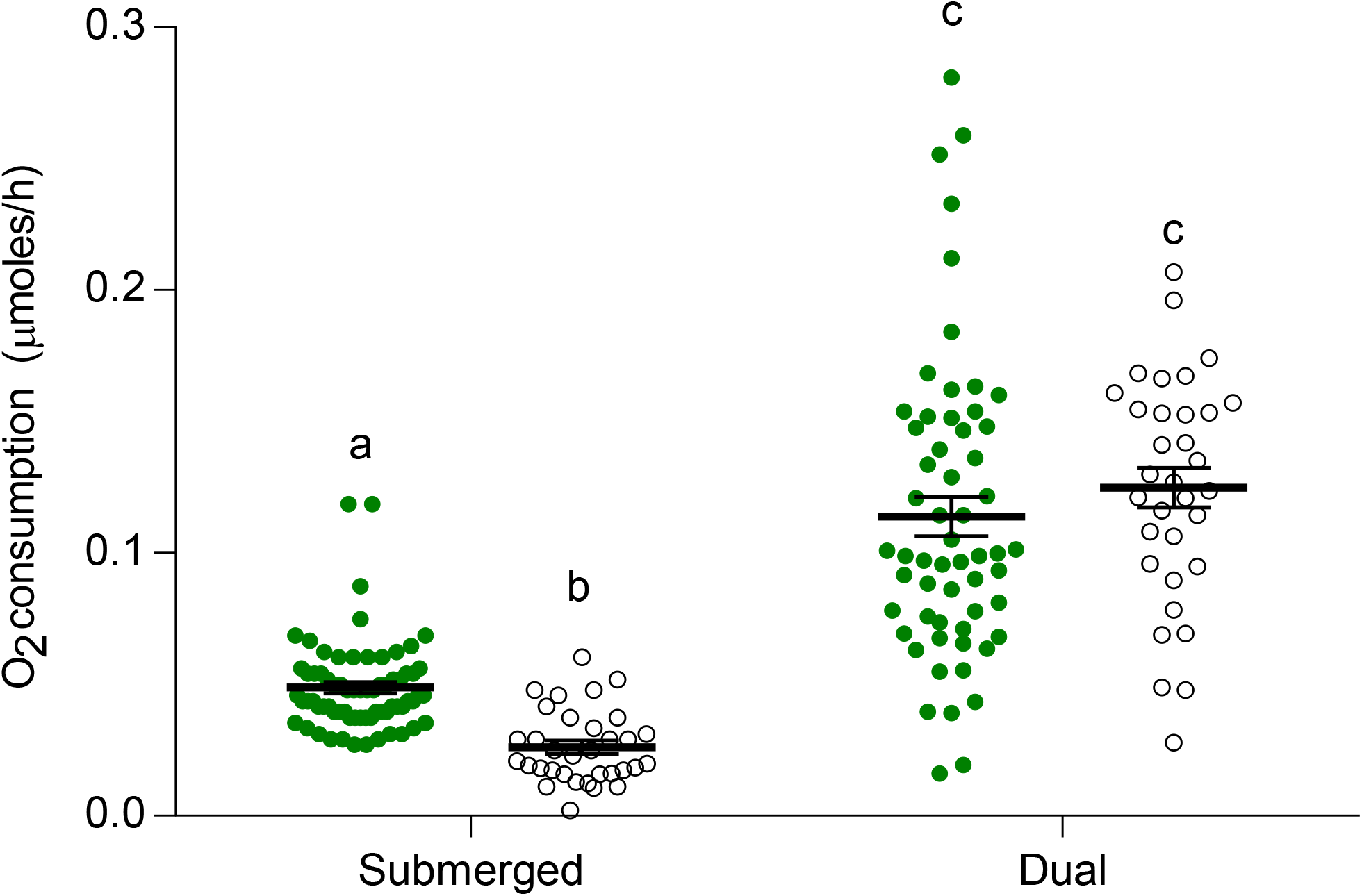
Total O_2_ consumption (mean and 95 % confidence intervals) of Ae. aegypti larvae and pupae with (dual) and without (submerged) access to air. Different letters indicate significant differences between treatments (Tukey multiple comparison).

#### Effect of temperature on oxygen consumption (Q_10_ of larvae)

The consumption of water-dissolved oxygen of *Ae. aegypti* larvae when they were submerged, also significantly varied with temperature (one-way ANOVA, F: 33.46, DF: 2, p<0.0001). The highest O_2_ consumption was registered at 35ºC, followed by 25ºC, and the lowest one was registered at 15ºC. The Q_10_ calculated between 15° and 25ºC was 1.47 and 1.66 between 25° and 35ºC, so the mean Q_10_ across the experiment was 1.56. Also, O_2_ consumption from air of *Ae. aegypti* larvae significantly varied with temperature (one-way ANOVA, F: 18.09, DF: 2, p<0.0001). The highest O_2_ consumption was registered at 35ºC, followed by 25ºC, and the lowest was registered at 15ºC. The Q_10_ calculated between 15° and 25ºC was 2.21 and between 25° and 35ºC was 1.45, so the mean Q_10_ across the experimental temperatures was 1.83.

#### Aedes albopictus

Similar patterns of O_2_ consumption were observed in *Ae. albopictus*. A significant difference in O_2_ consumption from the water was observed between vial conditions (one-way ANOVA, F: 278.36, DF: 1, p<0.0001, Fig. 5). *Ae. albopictus* larvae from the submerged condition treatment presented significantly higher O_2_ consumption than larvae from the closed vial (Fig. 5). In the closed vial condition, the mean dual (air + water) rate of O_2_ consumption was 0.0185 !moles of O_2_/h for larvae. They consumed a significantly higher amount of O_2_ (around 88%) from the air than from water (one-way ANOVA, F: 27.97, DF: 1, p< 0.0001, Fig. 6). Interestingly, no significant differences in the total O_2_ consumption of *Ae. albopictus* larvae were found between water-dissolved oxygen consumption when they were submerged or when they also had access to air (dual O_2_ consumption) in the closed vial condition (one-way ANOVA, F: 0.99, DF: 1, p= 0.325, Fig. 7).

**Figure 5.**
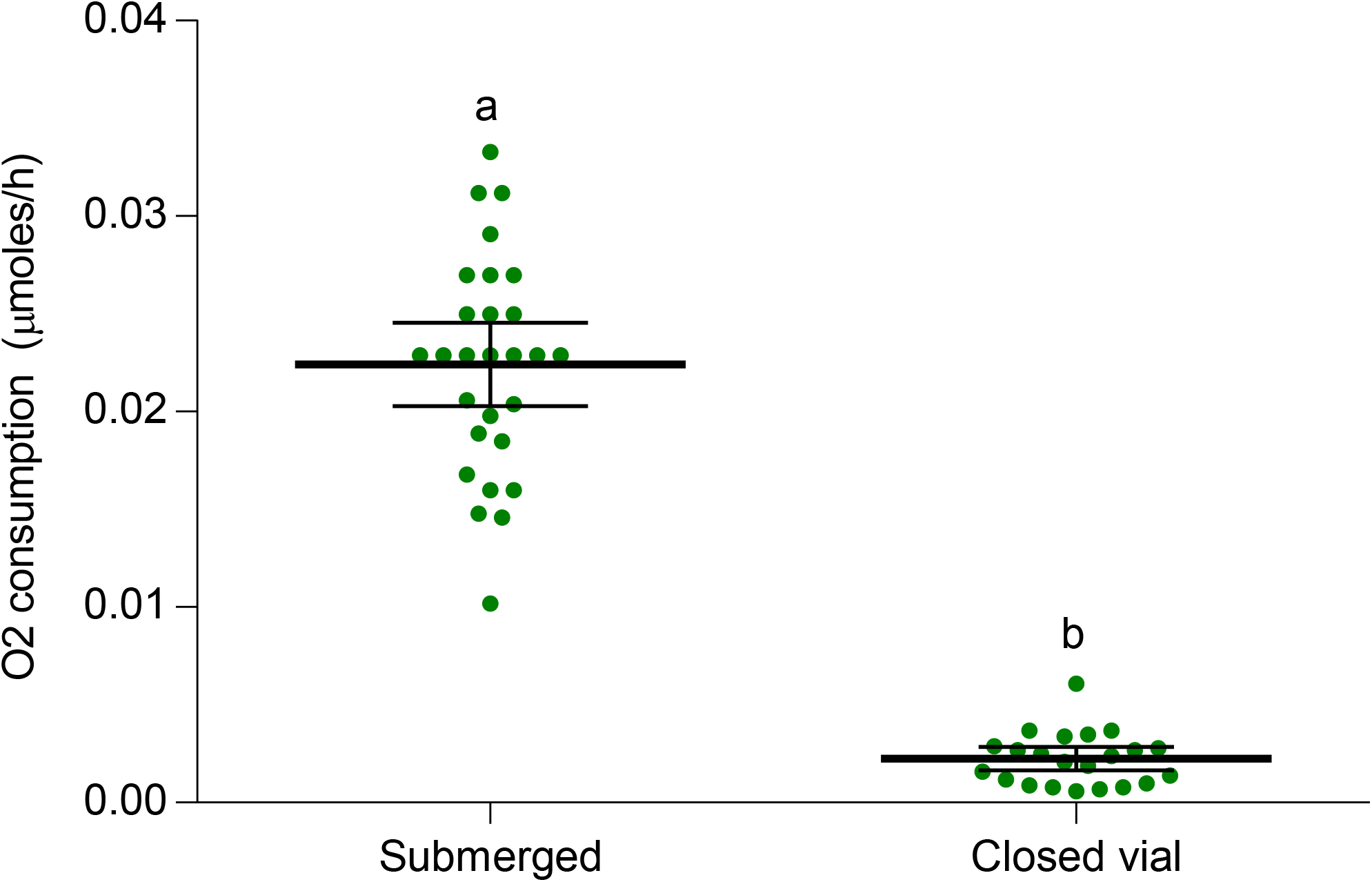
Consumption of O2 from water (mean and 95 % confidence intervals) by Ae. albopictus larvae under two conditions: submerged and closed vial. Different letters indicate significant differences between treatments.

**Figure 6.**
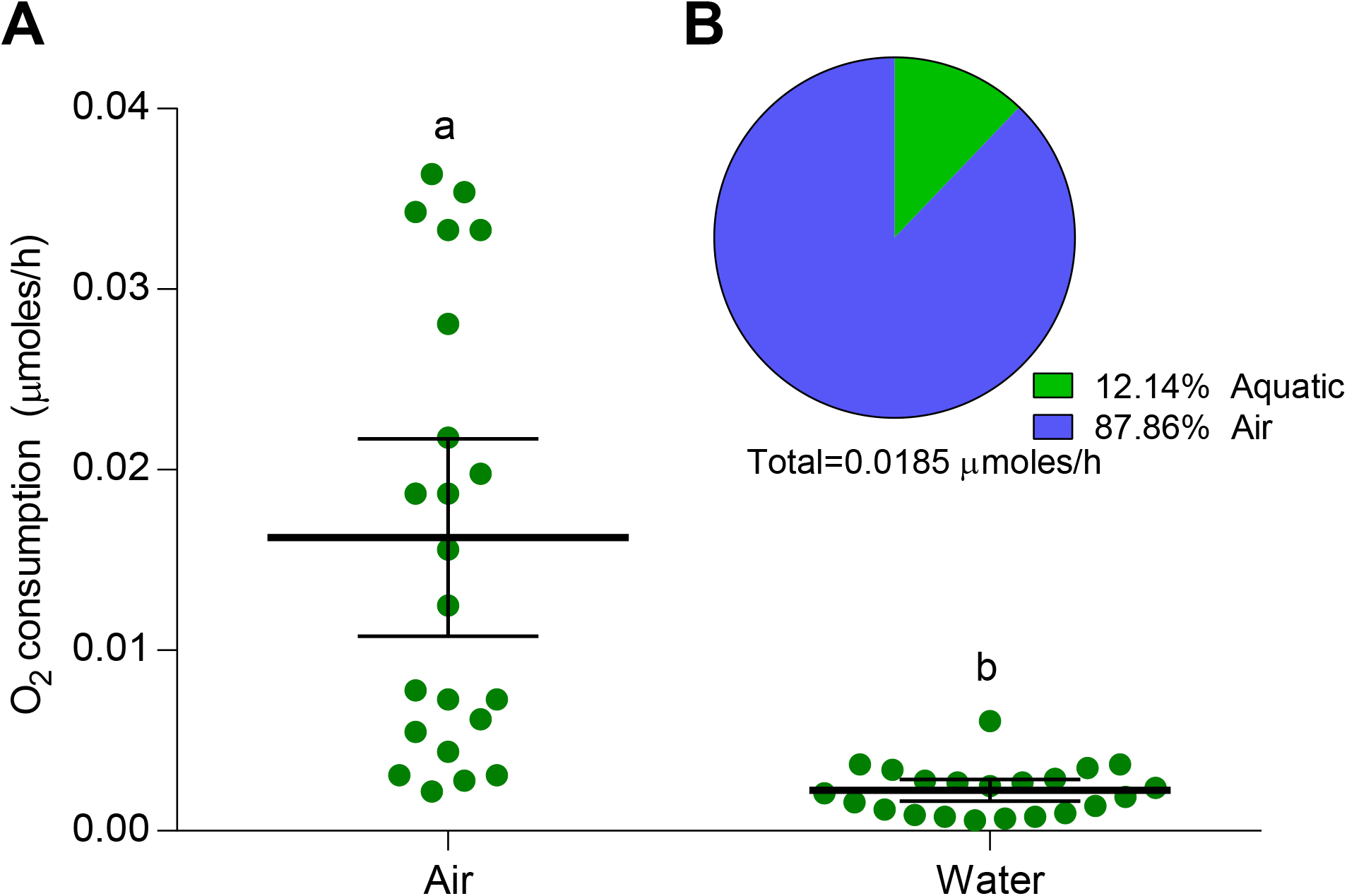
(**A**) O_2_ consumption (mean and 95 % confidence intervals) obtained from air and water of *Ae. albopictus* larvae in the closed vial condition. Percentage of oxygen consumption of *Ae. albopictus* larvae (**B**) obtained from air and water. Different letters indicate significant differences between treatments.

**Figure 7.**
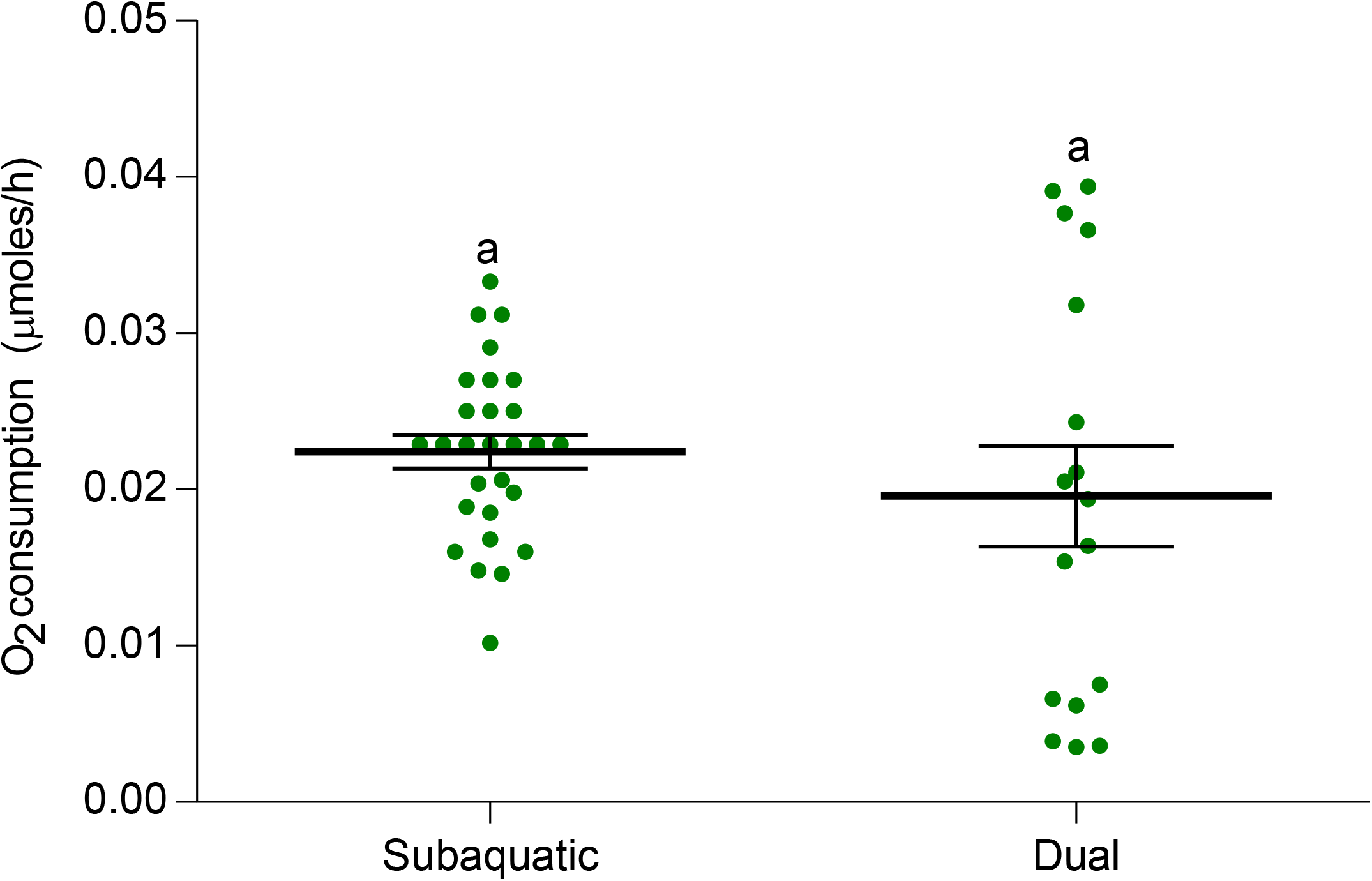
Total O_2_ consumption (mean and 95 % confidence intervals) of *Ae. albopictus* larvae with (dual) and without (submerged) access to air. Different letters indicate significant differences between treatments.

On the other hand, the temperature affected the rate of O_2_ consumption from water (one-way ANOVA, F: 77.52, DF: 2, p<0.0001). The rate of O_2_ consumption did not differ significantly between 15° and 25°C, but a significant increase was registered between either 15° or 25° and 35°C. The Q_10_ calculated between 15° and 25ºC was 1.47, and between 25° and 35ºC was 3.01, so the average Q_10_ between 15° and 35ºC was 2.24.

## Discussion

We report the first quantitative data on oxygen consumption, both from the air and from the water, by larvae and pupae of two major disease vectors, *Aedes aegypti* and *Ae. albopictus*. In both cases, far from a great surprise, most of the oxygen consumed comes from atmospheric air, but not all of it. The portion gathered from the water is low, but physiologically significant, which means it is just enough for survival, and much lower in pupae, which obtain practically the totality of the oxygen they consume from the air.

Even though underwater respiration by mosquito larvae has been repeatedly reported, this phenomenon remained anecdotal, deserving no attention by most people. In fact, not only there is no evidence to what extent oxygen gathered from water might be physiologically significant to mosquito larvae, but it has been completely ignored as a potential drawback in control procedures involving larval suffocation. Indeed, being detached from the surface and losing contact with the air, does not mean the immediate death of the larvae by asphyxiation. In complete submersion, they were capable of surviving for days, weeks, or even months, depending on the temperature of the water.

As can be expected for a poikilotherm and ectotherm organism, water temperature has a significant impact on larval metabolism, which is reflected in the intensity of both aerial and aquatic respiration. This dependence is quantitatively expressed by the calculation of Q_10_ values for oxygen consumption, and also by the differential survival time of completely submerged larvae across a wide range of temperatures. As expected, the lower the temperature, the longer the survival time observed. This fact can be explained by the modulation of larvae metabolism, reflected on the Q_10_ different from 1, together with a higher O_2_ concentration in the water, since the solubility of oxygen increases as temperature decreases. So, in tropical areas or areas where human activities result in relatively warm temperatures, we could expect a lower capacity to tolerate immersion for prolonged periods.

Interestingly, the moult cycle was markedly affected by the deprivation of access to atmospheric air. No larva was capable of accomplishing a normal moult in the prolonged submersion experiments. The oxygen gathered exclusively from the water resulted to be sufficient for surviving and swimming (larvae did not remain immobile nor in akinesis into the tubes), but not enough for moulting. Only larvae kept at the lower temperature showed signs of incomplete ecdysis after many weeks under water, which indicates that the first steps of the moult process took place. Probably, ecdysis is excessively expensive in terms of energetic demands to complete it under these conditions.

Aquatic respiration was reported several times in mosquito larvae. One of the first scientists to turn his interest towards water respiration in mosquito larvae was the Brazilian entomologist Ângelo Moreira Da Costa Lima more than a century ago. He performed a series of experiments placing larvae of different Culicidae species under complete submersion and reporting day by day the status of each individual (Da Costa Lima, 1914). The author observed that some larvae survived for several days and that one larva whose “leaflets” (i.e., anal papillae) had been removed, returned at the surface more often than another intact used as control. These results led the author to assert: *“The results of my experiments convinced me that mosquito larvae, while generally breathing mainly free air by the two tracheae of the respiratory syphon, also respire the oxygen of the air dissolved in water, the gaseous exchanges being made by the branchial leaflets and the general integument of the body.”*. This assertion was criticised by colleagues, who distrusted the results due to poor control of the experimental conditions (Sen, 1914). Da Costa Lima (1916) replicated some of the original experiments taking additional care and reporting similar observations to his previous ones. On the other hand, Da Costa Lima noted the resemblance to gills of anal papillae, in line with other colleagues considering these structures as respiratory organs in aquatic Diptera (Koch. 1938). The demonstration of the osmoregulatory function of the papillae (Wigglesworth, 1932, 1933, review by Bradley, 1987), together with the general assumption that the potential contribution of oxygen dissolved in the water should be insignificant, rapidly made aquatic respiration to be disregarded as physiologically relevant for mosquito larvae (Thorpe, 1933).

Other early investigations, such as those conducted by MacFie (1917), Ramsey and Carpenter (1932), Wang (1938), and Richards (1941) focussed on the oxygen requirements of mosquito larvae by submerging them in oxygenated or deoxygenated water, observing differential survival. According to our results (Fig. 5), and those obtained by Fraenkel and Herford (1938), fully submerged *Ae. aegypti* larvae would consume only half as much oxygen as larvae swimming in the normal manner. However, this is not the case in *Ae. albopictus*, whose larva can gather similar amounts of oxygen when fully submerged or attached to the surface (Fig. 8).

Using a different experimental approach, Krogh (1941) noticed that the gaseous pressure in the trachea of *Culex* larvae reduced during submersion; according to the author, this could be caused by the withdrawal of oxygen for respiration and the loss of CO_2_ through the external cuticle. This author estimated the volume of the tracheae close to 1.5 μlitre, suggesting that the content of oxygen would be enough for surviving 5-10 min underwater Krogh (1941).

Hagstrum (1970) measured the aerial and aquatic respiration of larvae of different species, in the presence of petroleum oils in the water. His study suggested that for *Ae. aegypti* aquatic respiration could represent *ca*. 5-20% of aerial respiration, rather than 50% as formerly proposed by Fraenkel and Herford (1938). This agrees with our results where the aquatic respiration of *Ae. aegypti* larvae constituted about 13%. Hagstrum (1970) also reported delayed mortality in *Ae. aegypti* larvae, although their tracheae were blocked with petroleum oil, noting that these larvae were unable to pupate.

Later on, different authors started to focus their attention on the fact that the amount of oxygen dissolved in the water might impact on the survival of larvae and pupae, and, as a consequence, eventually affect their control based on larval suffocation. Reiter (1978) kept submerged larvae of three mosquito species in water with fixed dissolved oxygen contents and at different temperatures. The author concluded that those larvicides which kill by anoxia are likely to be effective only when the water is less than 30% saturated with oxygen, the exact value depending on the species. Westwood et al. (1983) and Silberbush et al. (2015) extended this idea to larvae living in natural environments with access to the air, providing additional quantitative data related to survival and oxygen saturation in the water.

Interestingly, different authors commenced in recent years to insinuate that breathing in mosquito larvae may have been misunderstood, raising questions about canonical assumptions and the real effectiveness of larval suffocation as a method for controlling natural populations of mosquitoes. For instance, in an attempt to understand the underlying mechanism of breathing cut-off (e.g., wettability of the siphon), Lee et al. (2018) exposed larvae of *Aedes togoi* to water treated either with oil-film layers or with surfactants (i.e., substances impeding larvae to remain attached to the water surface). The survival times recorded were variable according to the treatment, but reached times of about one day. The authors concluded that cutaneous respiration dependent on the oxygen concentration in the water could affect the efficacy of larval asphyxiation methods. In this sense, Lee et al. (2018) underlined the need to prevent oxygen dissolution by blocking the exchange between the water and the atmospheric air at the surface. It is clear that this procedure would negatively affect not only mosquitoes, but also the rest of the organisms living in the same environment. More recently, Nyberg and Muto (2020) investigated the mechanism of action of what authors called *“mosquito acoustic larviciding”*. They reported that their acoustic treatment provoked the tracheal system a series of what they reported as *“previously unobserved phenomena”* According to Nyberg and Muto (2020), these phenomena would be difficult to explain based on the present knowledge of mosquito respiration, eventually concluding that the respiratory function in mosquitoes is far from being completely understood.

Our study sheds light on the respiratory physiology of the mosquito aquatic instars by analysing the survival and oxygen intake of mosquito larvae and pupae having or not having access to the air, and measuring the effect of the temperature on these variables. For the first time, we provide quantitative data on oxygen consumption from air and water, which have been measured separately and simultaneously. Our work also provides evidence demonstrating how our limited knowledge of crucial aspects of mosquito respiratory physiology may compromise control methods based on the suffocation of juvenile mosquitoes.

## Acknowledgements

Authors want to express their gratitude to the University of Tours, CNRS ANR and LESTUDIUM (France); CONICET, University of Buenos Aires and MINCYT (Argentina), The Company of Biologist (UK) and the ECOS-Sud programme (France-Argentina) A part of this work was conducted during the Master thesis of SG at the Univ. Tours. MSL is grateful to Nicolas and Simona Albanese, for their unconditional support.

## Supplementary material

**Fig S1.**
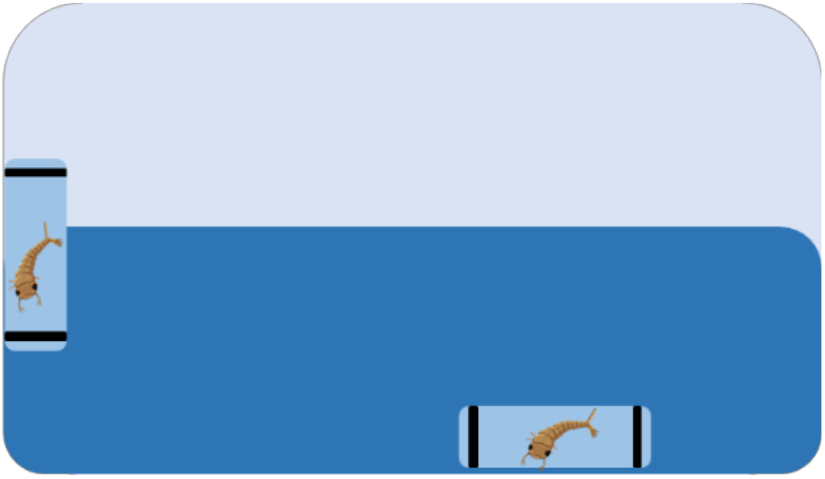
Setup for testing larval survival at different temperatures.

